# Identification of novel key biomarkers in Simpson-Golabi-Behmel Syndrome: Evidence from bioinformatics analysis

**DOI:** 10.1101/608927

**Authors:** Mujahed I. Mustafa, Abdelrahman H. Abdelmoneim, Nafisa M. Elfadol, Naseem S. Murshed, Zainab O. Mohammed, Mohamed A. Hassan

**Affiliations:** Department of Biotechnology, Africa city of Technology, Sudan; Hematology Department, Ribat University Hospital, Sudan

**Keywords:** Simpson-Golabi-Behmel Syndrome (SGBS), overgrowth Syndrome, *GPC3* bioinformatics analysis, SNPs, diagnostic markers

## Abstract

**Background:** The Simpson-Golabi-Behmel Syndrome (SGBS) or overgrowth Syndrome is a rare inherited X-linked condition characterized by pre- and postnatal overgrowth. The aim of the present study is to identify functional non-synonymous SNPs of GPC3 gene using various in silico approaches. These SNPs are supposed to have a direct effect on protein stability through conformation changes.

**Material and methods:** The SNPs were retrieved from the Single Nucleotide Polymorphism database (dbSNP) and further used to investigate a damaging effect using SIFT, PolyPhen, PROVEAN, SNAP2, SNPs&GO, PHD-SNP and P-mut, While we used I-mutant and MUPro to study the effect of SNPs on GPC3 protein structure. The 3D structure of human GPC3 protein is not available in the Protein Data Bank, so we used RaptorX to generate a 3D structural model for wild-type GPC3 to visualize the amino acids changes by UCSF Chimera. For biophysical validation we used project HOPE. Lastly we run conservational analysis by BioEdit and Consurf web server respectively.

**Results:** our results revealed three novel missense mutations (rs1460413167, rs1295603457 and rs757475450) that are found to be the most deleterious which effect on the GPC3 structure and function.

**Conclusion:** This present study could provide a novel insight into the molecular basis of overgrowth Syndrome.

## 1. Introduction

The Simpson-Golabi-Behmel Syndrome (SGBS) or overgrowth Syndrome is a rare inherited X-linked condition characterized by pre- and postnatal overgrowth, Affected individuals have several dysmorphisms that can include a distinct facial appearance, macroglossia, cleft palate, cardiac defects, enlarged and dysplastic kidneys, cryptorchidism, hypospadia, hernias, supernumerary nipples, vertebral and rib anomalies, coccygeal bony appendage, syndactyly, and polydactyly.[1–4] the first case was reported around 1940.[5] To date, two different clinical subtypes of SGBS have been defined. The typical SBGS (SGBS type I) [2–8] and a lethal and rare form, probably with less than 10 cases described known as SGBS type II. [9–11] In addition, these patients have an increased risk for the development of Wilms’ tumors.[12] Premature death is also very frequent.[13] Different mutations have been reported in SGBS type I.[14–22]

This dysplasia syndrome caused by loss-of-function mutations of the X-linked *GPC3* gene is localized on Xq26.1 [23, 24] which encodes a developmentally regulated cell membrane proteoglycan, glypican-3. [17, 19, 20, 25–29] that apparently plays a negative role in growth control by an unknown mechanism, However, outcomes from a detailed comparative analysis of growth patterns in dual mutants lacking GPC3 provided conclusive genetic evidence inconsistent with the theory that GPC3 performances as a growth suppressor by downregulating an IGF ligand.[29] Such a proteoglycan is inferred to play an important role in control and diagnosis in mesodermal tissues and in tumors predisposition.[30, 31] Some studies show association between *GPC3* gene and some types of human cancers.[32–36]

The aim of this study was to identify the pathogenic SNPs in the coding region of *GPC3* gene which causes SGBS type I using variant bioinformatics tools, Also to identify the most deleterious nsSNPs that could be used as diagnostic markers. A nsSNP is a single base change in a coding region that causes an amino acid change in its corresponding protein. If nsSNP modifies protein function, the change can have major phenotypic effects which are responsible for the pathology of the disease [37, 38] Genetic testing for mutations often reveals substitutions that are not easily categorized as pathogenic. Therefore a great effort has been done in translational bioinformatic tools for analysis of nsSNPs which have enhanced significantly in recent years and thus become more reliable for SNPs analysis.[39] Translational bioinformatics has become an important discipline in the era of personalized and precision medicine which aims to fill the gap between clinical and academic research by prioritizing the most pathogenic nsSNPs for further studies.[40–44] This is the first in silico analysis in the coding region of *GPC3* gene that prioritized nsSNPs for the larger population-based studies of overgrowth syndrome.

## 2. Material and methods

### 2.1 Data mining

The data on human *GPC3* gene was collected from National Center for Biological Information (NCBI) web site.[45] The SNP information (protein accession number and SNP ID) of the *GPC3* gene was retrieved from the NCBI dbSNP (http://www.ncbi.nlm.nih.gov/snp/) and the protein sequence was collected from Uniprot database. (https://www.uniprot.org/).

### 2.2 PolyPhen

PolyPhen (version 2) We used PolyPhen to study probable impacts of A.A. substitution on structural and functional properties of the protein by considering physical and comparative approaches.[46] It is available at (http://genetics.bwh.harvard.edu/pph2/).

### 2.3 PROVEAN

PROVEAN is a software tool which predicts whether an amino acid substitution or indel has an impact on the biological function of a protein. It is useful for filtering sequence variants to identify nonsynonymous or indel variants that are predicted to be functionally important.[47] It is available at (https://rostlab.org/services/snap2web/).

### 2.4 SNAP2

SNAP2 is a trained classifier that is based on a machine learning device called “neural network”. It distinguishes between effect and neutral variants/non-synonymous SNPs by taking a variety of sequence and variant features into account.[48] It is available at (https://rostlab.org/services/snap2web/).

### 2.5 SNPs&GO

SNP&GO is a method for the prediction of deleterious SNPs using protein functional annotation. The server is based on SVM classifier that discriminates between disease related and neutral SNPs. It has two components, one is sequence based and the other is structure based. The other method were used too (PHD-SNP and PANTHER).[49] It is available at (http://snps.biofold.org/snps-and-go/snps-and-go.html).

### 2.6 Stability Analysis

#### 2.6.1 I-Mutant 3.0

I-Mutant is an SVM-based tool for the automatic prediction of protein stability changes upon single point mutations. The predictions are performed starting either from the protein structure or, more importantly, from the protein sequence[50] It is available at (http://gpcr2.biocomp.unibo.it/cgi/predictors/I-Mutant3.0/IMutant3.0.cgi).

#### 2.6.2 MUpro

MUpro is a SVM-based tool for the prediction of protein stability changes upon nsSNPs. The value of the energy change is predicted, and a confidence score between −1 and 1 for measuring the confidence of the prediction is calculated. The accuracy for SVM using sequence information is 84.2%.[51] It is available at (http://mupro.proteomics.ics.uci.edu/).

### 2.7 Structural Analysis

#### 2.7.1 Modeling nsSNP locations on protein structure

Project hope is a new online web-server to search protein 3D structures (if available) by collecting structural information from a series of sources, including calculations on the 3D coordinates of the protein, sequence annotations from the UniProt database, and predictions by DAS services. Protein sequences were submitted to project hope server in order to analyze the structural and conformational variations that have resulted from single amino acid substitution corresponding to single nucleotide substitution. It is available at (http://www.cmbi.ru.nl/hope).

#### 2.7.2 Modeling Amino Acid Substitution

UCSF Chimera is a highly extensible program for interactive visualization and analysis of molecular structures and related data, including density maps, supramolecular assemblies, sequence alignments, docking results, conformational analysis Chimera (version 1.8).[52] It is available at (http://www.cgl.ucsf.edu/chimera/).

### 2.8 Conservational Analysis

#### 2.8.1 BioEdit

BioEdit software is intended to supply a single program that can handle most simple sequence and alignment editing and manipulation functions as well as a few basic sequences analyses. It is available for download at (http://www.mbio.ncsu.edu/bioedit/bioedit.html).

#### 2.8.2 Identification of Functional SNPs in Conserved Regions by using ConSurf server

ConSurf web server provides evolutionary conservation profiles for proteins of known structure in the PDB. Amino acid sequences similar to each sequence in the PDB were collected and multiply aligned using CSI-BLAST and MAFFT, respectively. The evolutionary conservation of each amino acid position in the alignment was calculated using the Rate 4Site algorithm, implemented in the ConSurf web server.[53] It is available at (http://consurf.tau.ac.il/).

## 3. Results

The effect of each SNP has been studied regarding function and stability of the protein using different in analysis tools with different considerations and features, in order to minimize the error to the least percentage possible. (Figure 1)

**Figure (1):**
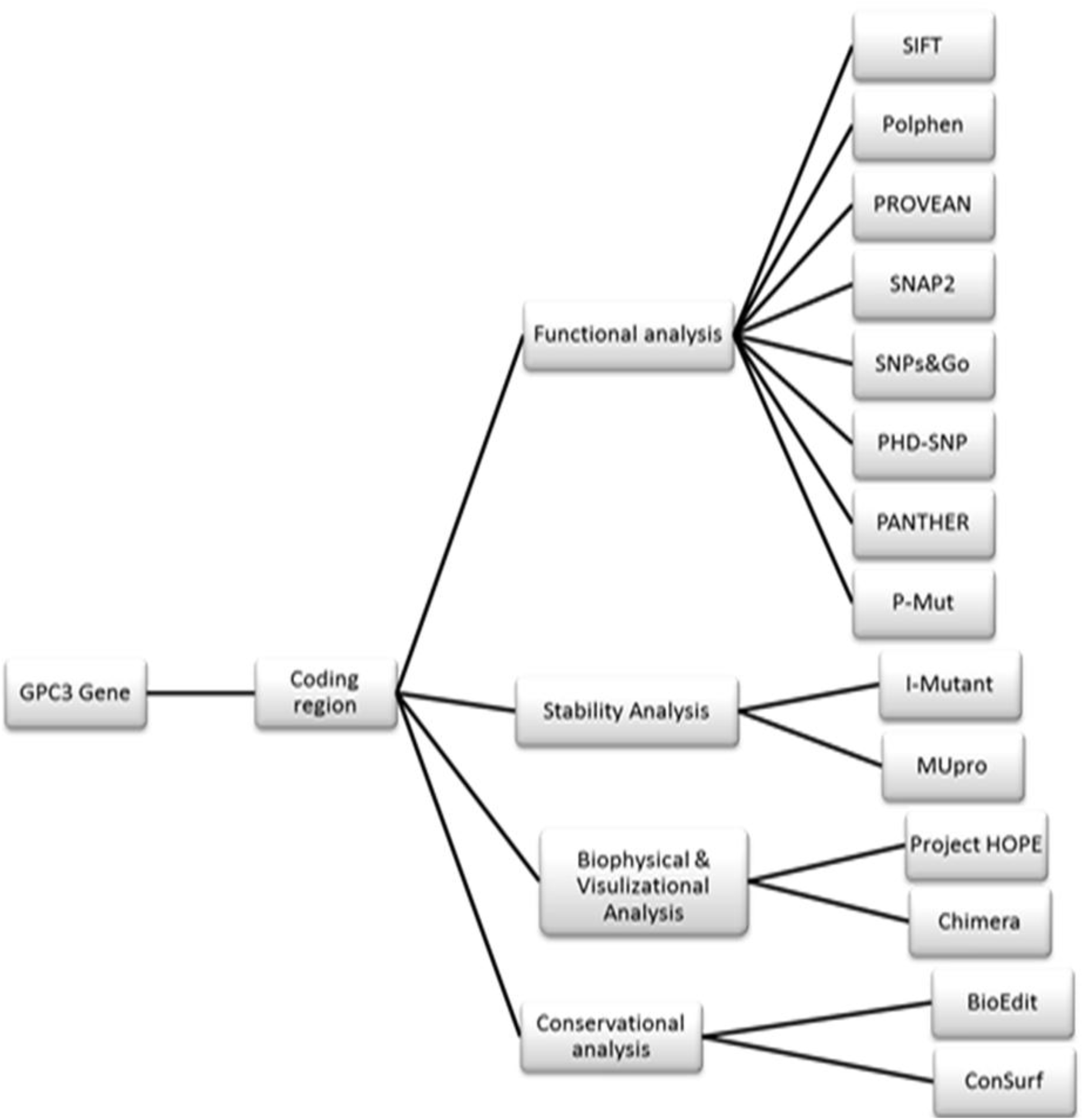
Diagrammatic Workflow Software’s used in SNPs analysis.

The total number of nsSNPs in the coding region of the *GPC3* gene in Human is 765 SNPs we retrieved from the dbSNP/NCBI database. There were 256 missense mutations then submitted them to functional analysis by SIFT, PolyPhen-2, PROVEAN and SNAP2 respectively. SIFT predicted 109 damaging SNPs, polyphen-2 predicted 115 damaging SNPs (50 possibly damaging and 65 probably damaging), PROVEAN predicted 82 damaging SNPs and SNAP2 predicted 127 deleterious SNPs. After filtering the Quadra-positive damaging SNPs, the number of SNPs reduced to 7 SNPs (Table 1) and we submitted them to SNP&GO, PANTHER, PhD-SNP and P-mut for further study their effect on the function. PhD-SNP and P-mut predicted 7 disease associated SNPs, SNP&GO predicted 5, and while PANTHER predicted 5 disease associated SNPs. So we filtered the Quadra-positive disease associated SNPs the number reduce to 4 SNPs (Table 2) and submitted them to I-Mutant3.0 and MUPro to study their effect on the stability. All the SNPs were found to cause a decrease in the stability of the protein except for one SNP (G257D) predicted by I-Mutant 3.0 to increase the stability. (Table 3)

**Table (1):** Damaging or Deleterious nsSNPs associated predicted by various softwares:

| dbSNP rs# | sub | SIFT<br>Prediction | Score | Polyphen Prediction | Score | PROVEAN<br>Prediction | Score | SNAP2<br>Prediction | Score |
| --- | --- | --- | --- | --- | --- | --- | --- | --- | --- |
| - | D500Y | Affect | 0 | probably damaging | 1 | Deleterious | -4.572 | effect | 68 |
| rs1203009272 | G440R | Affect | 0 | probably damaging | 1 | Deleterious | -5.147 | effect | 90 |
| rs104894854 | W296R | Affect | 0 | probably damaging | 1 | Deleterious | -13.836 | effect | 98 |
| rs1460413167 | P212H | Affect | 0 | probably damaging | 1 | Deleterious | -7.77 | effect | 73 |
| rs140848049 | F208L | Affect | 0 | probably damaging | 1 | Deleterious | -5.913 | effect | 78 |
| rs1295603457 | C65Y | Affect | 0 | probably damaging | 1 | Deleterious | -7.572 | effect | 87 |
| rs757475450 | R39C | Affect | 0 | probably damaging | 1 | Deleterious | -4.575 | effect | 59 |

**Table (2):** Disease effect nsSNPs associated variations predicted by predicted by various softwares:

| sub | SNPs&GO Prediction | RI | Probability | PANTHER Prediction | RI | Probability | PHD Prediction | RI | Probability | P-mut Prediction | Probability |
| --- | --- | --- | --- | --- | --- | --- | --- | --- | --- | --- | --- |
| W296R | Disease | 7 | 0.858 | Disease | 10 | 0.99 | Disease | 9 | 0.925 | Disease | 0.89 (92%) |
| P212H | Disease | 4 | 0.709 | Disease | 10 | 0.994 | Disease | 6 | 0.788 | Disease | 0.81 (89%) |
| C65Y | Disease | 6 | 0.787 | Disease | 8 | 0.878 | Disease | 7 | 0.85 | Disease | 0.80 (89%) |
| R39C | Disease | 3 | 0.63 | Disease | 8 | 0.903 | Disease | 4 | 0.687 | Disease | 0.73 (87%) |

**\*RI:** Reliability Index

**Table (3):** stability analysis predicted by I-Mutant version 3.0 and MUPro:

**\*RI:** Reliability Index **Table (3):** stability analysis predicted by I-Mutant version 3.0 and MUPro:
| dbSNP rs# | Sub | SVM2 Prediction Effect | RI | DDG Value Prediction | Mupro Prediction | Mupro Score |
| --- | --- | --- | --- | --- | --- | --- |
| rs104894854 | W296R | Decrease | 8 | -0.99 | Decrease | -0.95856 |
| rs1460413167 | P212H | Decrease | 8 | -1.67 | Decrease | -1.30249 |
| rs1295603457 | C65Y | Increase | 0 | -0.09 | Decrease | -0.56309 |
| rs757475450 | R39C | Decrease | 3 | -0.74 | Decrease | -0.28424 |

## 4. Discussion

Three novel mutations were found to have a damaging effect on the stability and function of the *GPC3* gene using bioinformatics analysis.

There is a study that has been reported which shows a missense mutation that causes SGBS, [19] which matches with this study findings. Some studies show association between *GPC3* gene and some types of liver cancer such as hepatocellular carcinoma.[30, 34, 54, 55] Therefore, this study can open the door for novel diagnostic biomarkers for hepatocellular carcinoma. Combination detection of serum GPC3 and pathogenic SNPs through clinical and genetic testing must be positively matched; this can enhance accuracy and efficiency of hepatocellular carcinoma diagnosis.

In addition it confirms that (W296R) is pathogenic; this result shows similarities with the result found earlier in dbSNPs/NCBI database. Furthermore, all these SNPs (P212H, C65Y, R39C) were retrieved as untested, in this study were found to be all pathogenic.

The most four deleterious SNPs were submitted to project HOPE which revealed that all they are located in a domain in the protein and thus might have a dynamic change in the structure and function of the protein and it may affect its ability to bind with its targets, in (R39C): The wild-type residue charge was positive, the mutant residue charge is neutral; Due to the loss of charge of the wild-type residue, this can cause loss of interactions with other molecules or residues. The mutant residue is more hydrophobic than the wild-type residue; in arrears of this change, this can result in loss of hydrogen bonds and/or disturb correct folding. In (C65Y): The wild-type residue is annotated in UniProt to be involved in a cysteine bridge, which is important for stability of the protein. Only cysteines can make these type of bonds, the mutation causes loss of this interaction and will have a severe effect on the 3D-structure of the protein. In (P212H): The wild-type residue is a proline. Prolines are known to be very rigid and therefore induce a special backbone conformation which might be required at this position. The mutation can disturb this special conformation. While in (W296R): This mutation is located in a region with known splice variants and matches a previously described variant, with the following description: Simpson-Golabi-Behmel syndrome 1 (SGBS1) [MIM: 312870]. The hydrophobicity of the wild-type and mutant residue differs; hydrophobic interactions, either in the core of the protein or on the surface, will be lost. While UCSF Chimera were used to visualize the amino acids change. (figures 2–5)

**Figure (2):**
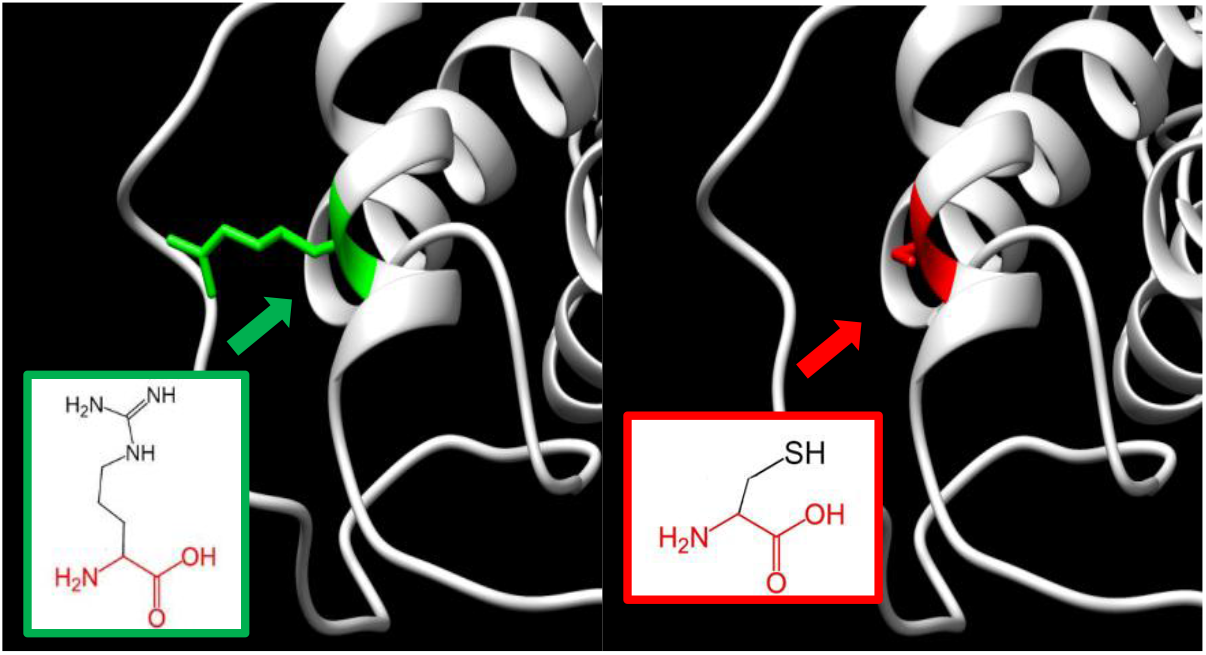
(rs757475450) (R39C) the amino acid Arginine changes to Cysteine at position 39. Illustrated by chimera (v 1.8) and project HOPE.

**Figure (3):**
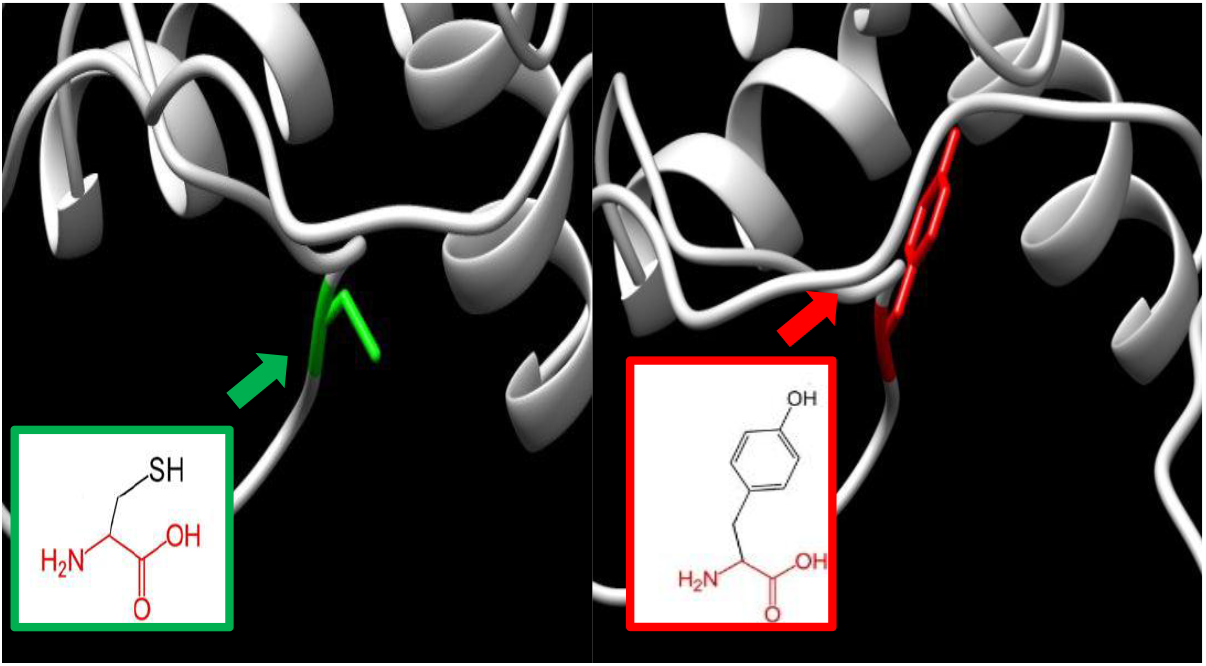
(rs1295603457) (C65Y) the amino acid Cysteine changes to Tyrosine at position 65. Illustrated by chimera (v 1.8) and project HOPE.

**Figure (4):**
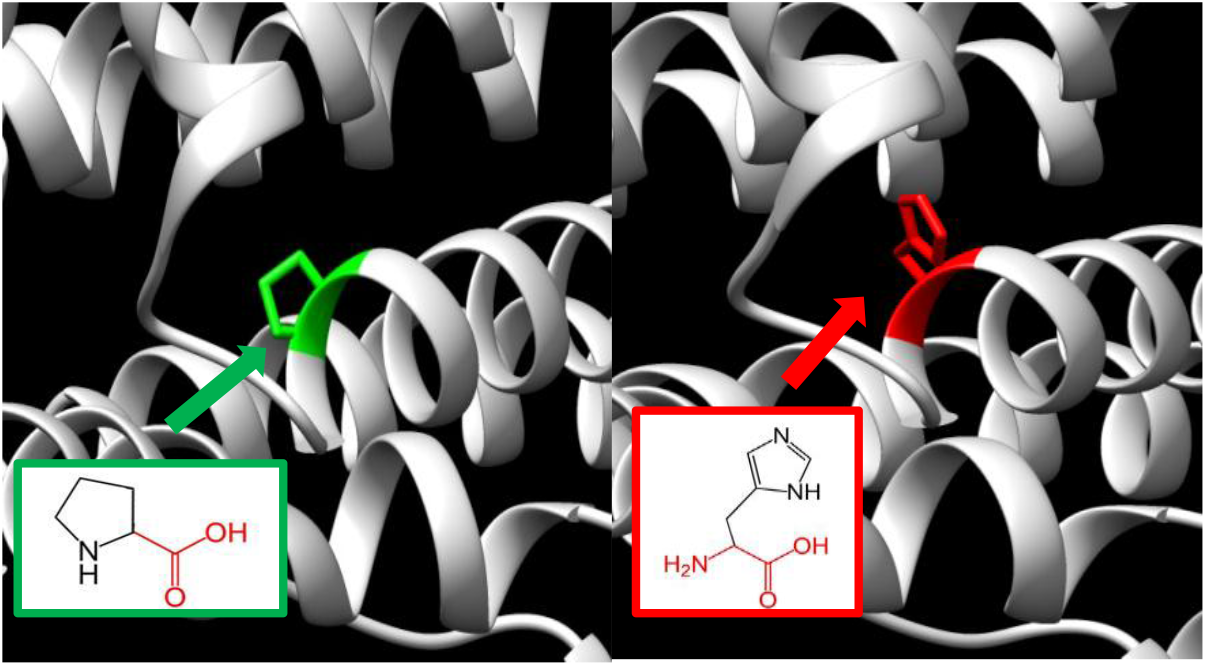
(rs1460413167) (P212H) the amino acid Proline changes to Histidine at position 65. Illustrated by chimera (v 1.8) and project HOPE.

**Figure (5):**
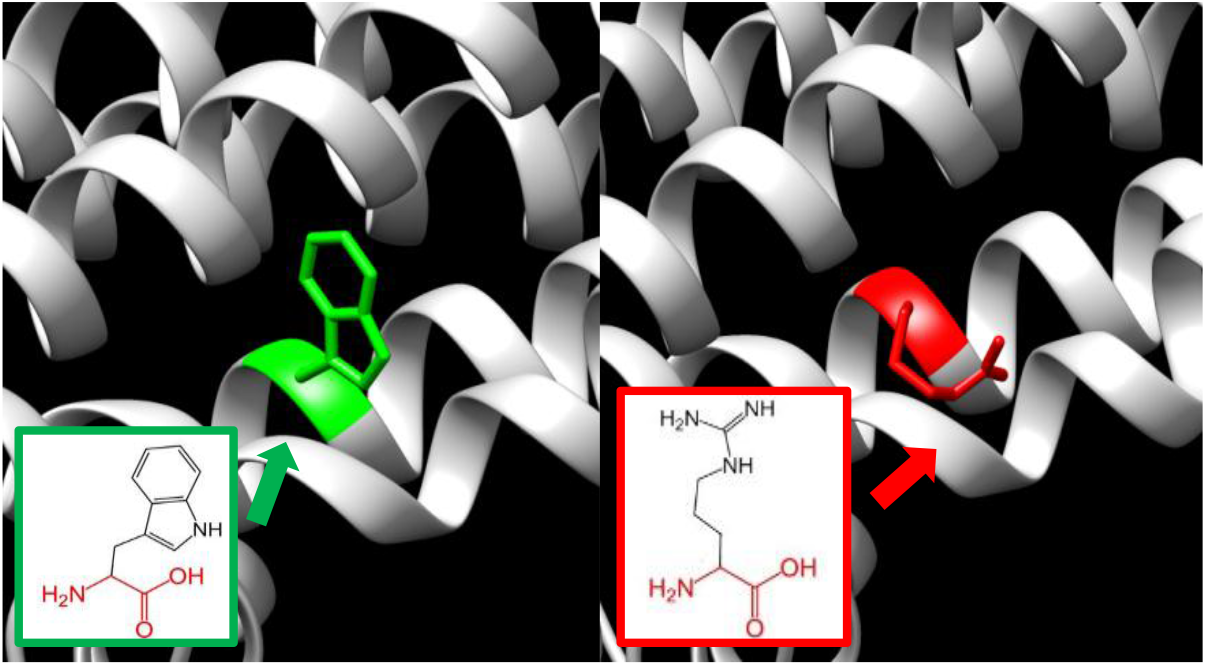
(rs104894854) (W296R) the amino acid Tryptophan changes to Arginine at position 296. Illustrated by chimera (v1.8) and project HOPE.

We also observed that, all the four SNPs were found in conserve region. (Figure 6) The seven amino acid sequences of GPC3 were retrieved from UniProt database.[56] While Sequence Alignments were done by BioEdit (v7.2.5). We also used ConSurf server; the nsSNPs that are located at highly conserved amino acid positions have a tendency to be the most damaging nsSNPs. (Figure 7)

**Figure (6):**
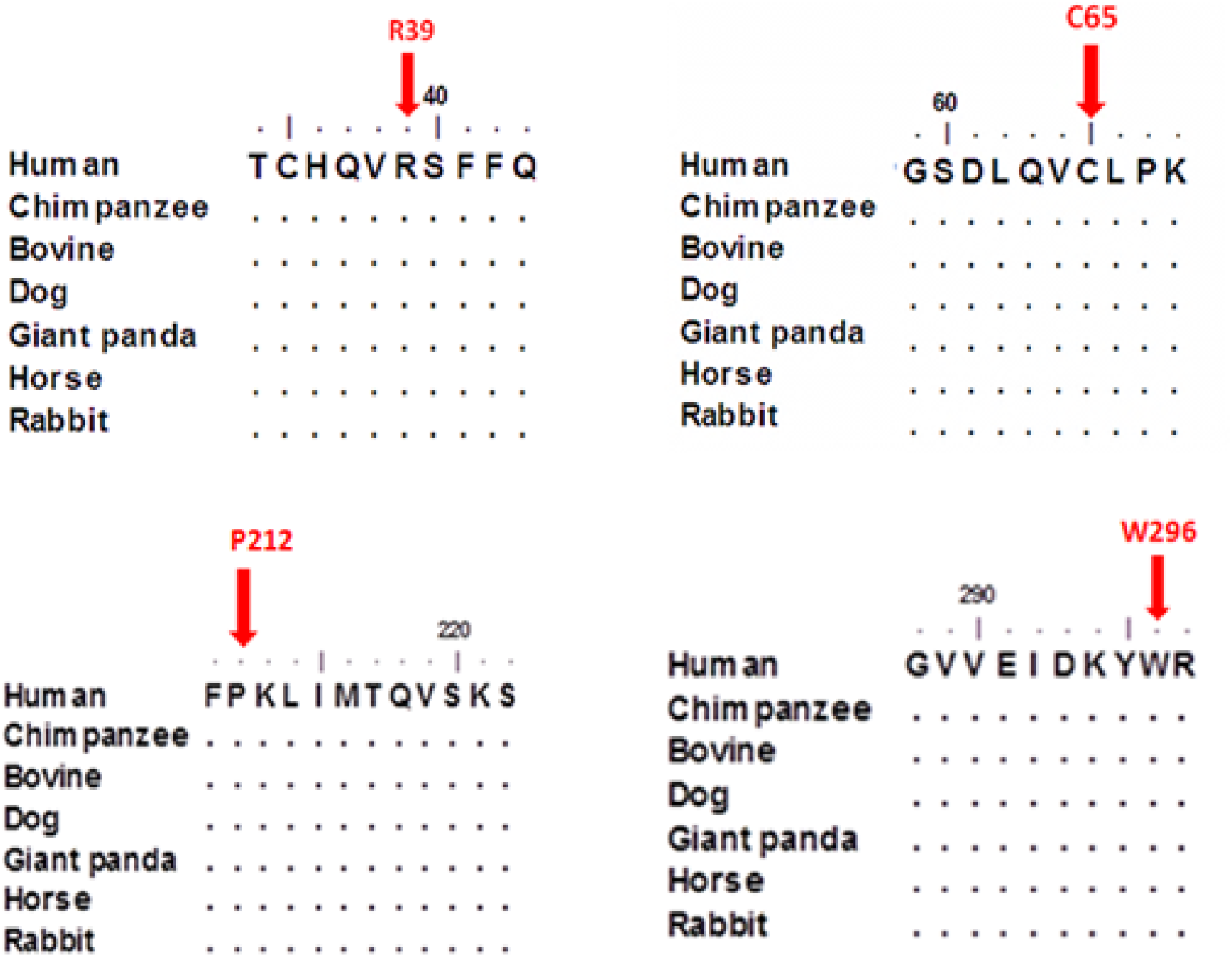
*GPC3* Family of seven amino acid sequences demonstrating that the residues predicted to be mutated (indicated by red arrows) are evolutionarily conserved across species. Sequences Alignment was done by BioEdit (v7.2.5).

**Figure (7):**
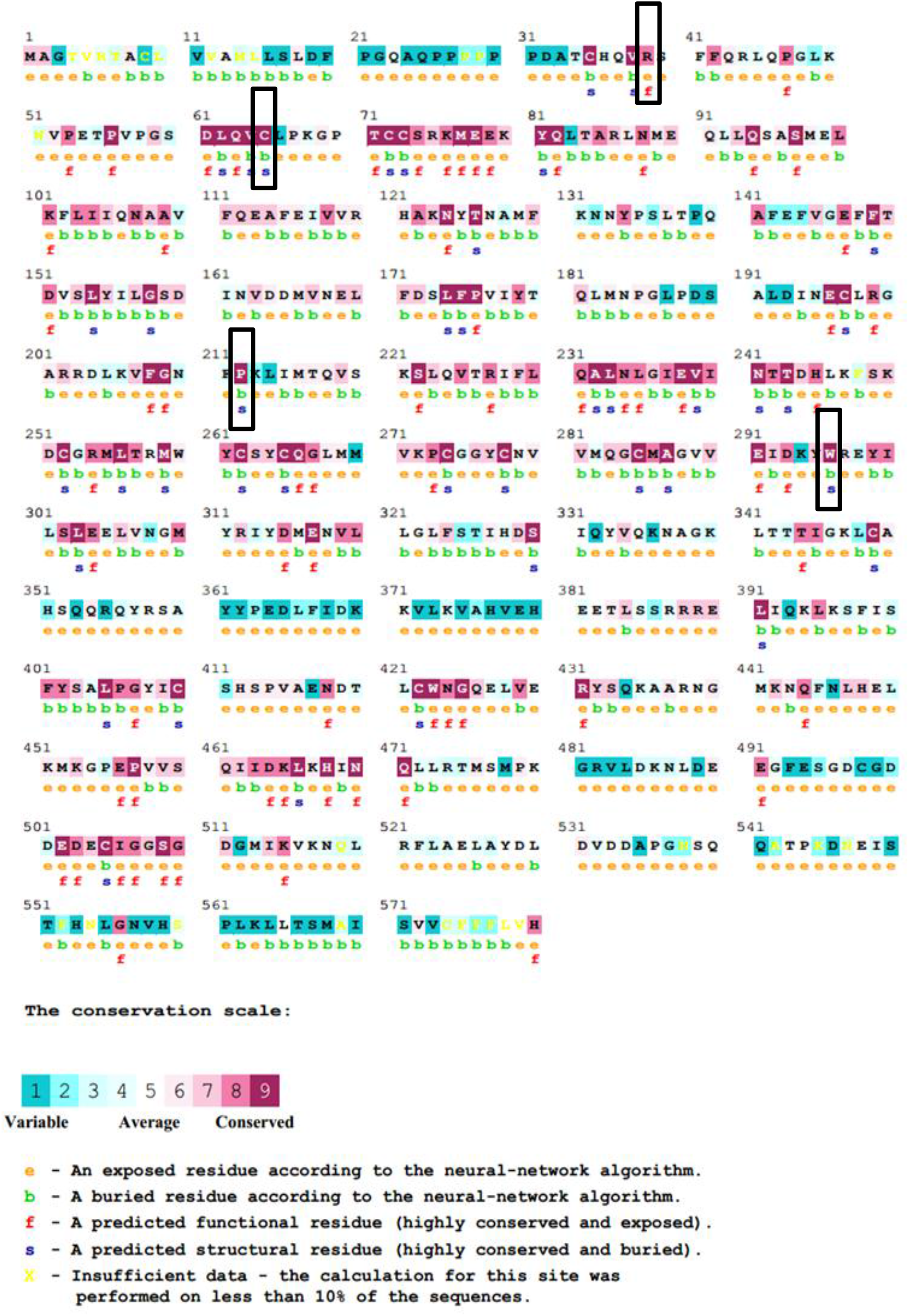
Unique and conserver regions in GPC3 protein were determined using Consurf web server.

This study is the first in analysis approach while all other previous studies were in vitro, in vivo and whole exome sequencing. [57–60] It revealed three novel missense mutations that have a potential functional impact and may thus be used as diagnostic markers for overgrowth syndrome.[61] Finally some appreciations of wet lab techniques are recommended to support these findings.

## 5. Conclusion

The present study could provide a novel insight into the molecular basis of overgrowth Syndrome by evidence from bioinformatics analysis. Three novel missense mutations (rs757475450, rs1295603457 and rs1460413167) that have a potential functional impact and may consequently can be used as diagnostic markers for overgrowth syndrome. As well as these SNPs can be used for the larger population-based studies of overgrowth syndrome.

## Acknowledgment

The authors wish to acknowledgment the enthusiastic cooperation of Africa City of Technology - Sudan.

## References

[1] E. Cottereau, I. Mortemousque, M. P. Moizard, L. Burglen, D. Lacombe, B. Gilbert-Dussardier, et al., “Phenotypic spectrum of Simpson-Golabi-Behmel syndrome in a series of 42 cases with a mutation in GPC3 and review of the literature,” Am J Med Genet C Semin Med Genet, vol. 163c, pp. 92–105, May 2013.

[2] A. Behmel, E. Plochl, and W. Rosenkranz, “A new X-linked dysplasia gigantism syndrome: identical with the Simpson dysplasia syndrome?,” Hum Genet, vol. 67, pp. 409–13, 1984.

[3] M. Golabi and L. Rosen, “A new X-linked mental retardation-overgrowth syndrome,” Am J Med Genet, vol. 17, pp. 345–58, Jan 1984.

[4] M. R. DeBaun, J. Ess, and S. Saunders, “Simpson Golabi Behmel syndrome: progress toward understanding the molecular basis for overgrowth, malformation, and cancer predisposition,” Mol Genet Metab, vol. 72, pp. 279–86, Apr 2001.

[5] F. Gurrieri, M. G. Pomponi, R. Pietrobono, E. Lucci-Cordisco, E. Silvestri, G. Storniello, et al., “The Simpson-Golabi-Behmel syndrome: A clinical case and a detective story,” Am J Med Genet A, vol. 155a, pp. 145–8, Jan 2011.

[6] J. L. Simpson, S. Landey, M. New, and J. German, “A previously unrecognized X-linked syndrome of dysmorphia,” Birth Defects Orig Artic Ser, vol. 11, pp. 18–24, 1975.

[7] M. L. Vuillaume, M. P. Moizard, S. Rossignol, E. Cottereau, S. Vonwill, J. L. Alessandri, et al., “Mutation update for the GPC3 gene involved in Simpson-Golabi-Behmel syndrome and review of the literature,” Hum Mutat, vol. 39, pp. 2110–2112, Dec 2018.

[8] G. Neri and M. Moscarda, “Overgrowth syndromes: a classification,” Endocr Dev, vol. 14, pp. 53–60, 2009.

[9] J. Tenorio, P. Arias, V. Martinez-Glez, F. Santos, S. Garcia-Minaur, J. Nevado, et al., “Simpson-Golabi-Behmel syndrome types I and II,” Orphanet J Rare Dis, vol. 9, p. 138, Sep 20 2014.

[10] L. M. Brzustowicz, S. Farrell, M. B. Khan, and R. Weksberg, “Mapping of a new SGBS locus to chromosome Xp22 in a family with a severe form of Simpson-Golabi-Behmel syndrome,” Am J Hum Genet, vol. 65, pp. 779–83, Sep 1999.

[11] D. Terespolsky, S. A. Farrell, J. Siegel-Bartelt, and R. Weksberg, “Infantile lethal variant of Simpson-Golabi-Behmel syndrome associated with hydrops fetalis,” Am J Med Genet, vol. 59, pp. 329–33, Nov 20 1995.

[12] R. M. Hughes-Benzie, A. G. Hunter, J. E. Allanson, and A. E. Mackenzie, “Simpson-Golabi-Behmel syndrome associated with renal dysplasia and embryonal tumor: localization of the gene to Xqcen-q21,” Am J Med Genet, vol. 43, pp. 428–35, Apr 15-May 1 1992.

[13] C. L. Garganta and J. N. Bodurtha, “Report of another family with Simpson-Golabi-Behmel syndrome and a review of the literature,” Am J Med Genet, vol. 44, pp. 129–35, Sep 15 1992.

[14] C. B. Griffith, R. C. Probert, and G. H. Vance, “Genital anomalies in three male siblings with Simpson-Golabi-Behmel syndrome,” Am J Med Genet A, vol. 149a, pp. 2484–8, Nov 2009.

[15] A. Vaisfeld, M. G. Pomponi, R. Pietrobono, E. Tabolacci, and G. Neri, “Simpson-Golabi-Behmel syndrome in a female: A case report and an unsolved issue,” Am J Med Genet A, vol. 173, pp. 285–288, Jan 2017.

[16] S. Halayem, M. Hamza, F. Maazoul, H. Ben Turkia, M. Touati, N. Tebib, et al., “Distinctive findings in a boy with Simpson-Golabi-Behmel syndrome,” Am J Med Genet A, vol. 170a, pp. 1035–9, Apr 2016.

[17] M. Li, C. Shuman, Y. L. Fei, E. Cutiongco, H. A. Bender, C. Stevens, et al., “GPC3 mutation analysis in a spectrum of patients with overgrowth expands the phenotype of Simpson-Golabi-Behmel syndrome,” Am J Med Genet, vol. 102, pp. 161–8, Aug 1 2001.

[18] S. Mariani, L. Iughetti, R. Bertorelli, D. Coviello, M. Pellegrini, A. Forabosco, et al., “Genotype/phenotype correlations of males affected by Simpson-Golabi-Behmel syndrome with GPC3 gene mutations: patient report and review of the literature,” J Pediatr Endocrinol Metab, vol. 16, pp. 225–32, Feb 2003.

[19] M. Veugelers, B. D. Cat, S. Y. Muyldermans, G. Reekmans, N. Delande, S. Frints, et al., “Mutational analysis of the GPC3/GPC4 glypican gene cluster on Xq26 in patients with Simpson-Golabi-Behmel syndrome: identification of loss-of-function mutations in the GPC3 gene,” Hum Mol Genet, vol. 9, pp. 1321–8, May 22 2000.

[20] G. Pilia, R. M. Hughes-Benzie, A. MacKenzie, P. Baybayan, E. Y. Chen, R. Huber, et al., “Mutations in GPC3, a glypican gene, cause the Simpson-Golabi-Behmel overgrowth syndrome,” Nat Genet, vol. 12, pp. 241–7, Mar 1996.

[21] K. Ridnoi, E. Kurvinen, S. Pajusalu, T. Reimand, and K. Ounap, “Two Consecutive Pregnancies with Simpson-Golabi-Behmel Syndrome Type 1: Case Report and Review of Published Prenatal Cases,” Mol Syndromol, vol. 9, pp. 205–213, Jul 2018.

[22] H. K. Stove, N. Becher, V. Gjorup, M. Ramsing, I. Vogel, and E. M. Vestergaard, “First reported case of Simpson-Golabi-Behmel syndrome in a female fetus diagnosed prenatally with chromosomal microarray,” Clin Case Rep, vol. 5, pp. 608–612, May 2017.

[23] G. Jedraszak, M. Girard, A. Mellos, D. D. Djeddi, C. Chardot, A. Vanrenterghem, et al., “A patient with Simpson-Golabi-Behmel syndrome, biliary cirrhosis and successful liver transplantation,” Am J Med Genet A, vol. 164a, pp. 774–7, Mar 2014.

[24] R. Kosaki, T. Takenouchi, N. Takeda, M. Kagami, K. Nakabayashi, K. Hata, et al., “Somatic CTNNB1 mutation in hepatoblastoma from a patient with Simpson-Golabi-Behmel syndrome and germline GPC3 mutation,” Am J Med Genet A, vol. 164a, pp. 993–7, Apr 2014.

[25] R. Huber, L. Crisponi, R. Mazzarella, C. N. Chen, Y. Su, H. Shizuya, et al., “Analysis of exon/intron structure and 400 kb of genomic sequence surrounding the 5’-promoter and 3’-terminal ends of the human glypican 3 (GPC3) gene,” Genomics, vol. 45, pp. 48–58, Oct 1 1997.

[26] J. Schmidt, R. Hollstein, F. J. Kaiser, and G. Gillessen-Kaesbach, “Molecular analysis of a novel intragenic deletion in GPC3 in three cousins with Simpson-Golabi-Behmel syndrome,” Am J Med Genet A, vol. 173, pp. 1400–1405, May 2017.

[27] D. D. Villarreal, H. Villarreal, A. M. Paez, D. Peppas, J. Lynch, E. Roeder, et al., “A patient with a unique frameshift mutation in GPC3, causing Simpson-Golabi-Behmel syndrome, presenting with craniosynostosis, penoscrotal hypospadias, and a large prostatic utricle,” Am J Med Genet A, vol. 161a, pp. 3121–5, Dec 2013.

[28] S. Yano, B. Baskin, A. Bagheri, Y. Watanabe, K. Moseley, A. Nishimura, et al., “Familial Simpson-Golabi-Behmel syndrome: studies of X-chromosome inactivation and clinical phenotypes in two female individuals with GPC3 mutations,” Clin Genet, vol. 80, pp. 466–71, Nov 2011.

[29] E. Chiao, P. Fisher, L. Crisponi, M. Deiana, I. Dragatsis, D. Schlessinger, et al., “Overgrowth of a mouse model of the Simpson-Golabi-Behmel syndrome is independent of IGF signaling,” Dev Biol, vol. 243, pp. 185–206, Mar 1 2002.

[30] P. Lapunzina, I. Badia, C. Galoppo, E. De Matteo, P. Silberman, A. Tello, et al., “A patient with Simpson-Golabi-Behmel syndrome and hepatocellular carcinoma,” J Med Genet, vol. 35, pp. 153–6, Feb 1998.

[31] R. Savarirayan and A. Bankier, “Simpson-Golabi-Behmel syndrome and attention deficit hyperactivity disorder in two brothers,” J Med Genet, vol. 36, pp. 574–6, Jul 1999.

[32] W. X. Bai, J. Gao, C. Qian, and X. Q. Zhang, “[A bioinformatics analysis of differentially expressed genes associated with liver cancer],” Zhonghua Gan Zang Bing Za Zhi, vol. 25, pp. 435–439, Jun 20 2017.

[33] F. Cartier, E. Indersie, S. Lesjean, J. Charpentier, K. B. Hooks, A. Ghousein, et al., “New tumor suppressor microRNAs target glypican-3 in human liver cancer,” Oncotarget, vol. 8, pp. 41211–41226, Jun 20 2017.

[34] Y. Wu, H. Liu, and H. Ding, “GPC-3 in hepatocellular carcinoma: current perspectives,” J Hepatocell Carcinoma, vol. 3, pp. 63–67, 2016.

[35] J. Filmus, “Glypicans in growth control and cancer,” Glycobiology, vol. 11, pp. 19r–23r, Mar 2001.

[36] H. Lin, R. Huber, D. Schlessinger, and P. J. Morin, “Frequent silencing of the GPC3 gene in ovarian cancer cell lines,” Cancer Res, vol. 59, pp. 807–10, Feb 15 1999.

[37] G. Shaw, “Polymorphism and single nucleotide polymorphisms (SNPs),” BJU Int, vol. 112, pp. 664–5, Sep 2013.

[38] Y. Chen, F. Cunningham, D. Rios, W. M. McLaren, J. Smith, B. Pritchard, et al., “Ensembl variation resources,” BMC Genomics, vol. 11, p. 293, May 11 2010.

[39] C. George Priya Doss, C. Sudandiradoss, R. Rajasekaran, P. Choudhury, P. Sinha, P. Hota, et al., “Applications of computational algorithm tools to identify functional SNPs,” Funct Integr Genomics, vol. 8, pp. 309–16, Nov 2008.

[40] J. D. Tenenbaum, “Translational Bioinformatics: Past, Present, and Future,” Genomics Proteomics Bioinformatics, vol. 14, pp. 31–41, Feb 2016.

[41] J. Vamathevan and E. Birney, “A Review of Recent Advances in Translational Bioinformatics: Bridges from Biology to Medicine,” Yearb Med Inform, vol. 26, pp. 178–187, Aug 2017.

[42] P. Katara, “Single nucleotide polymorphism and its dynamics for pharmacogenomics,” Interdiscip Sci, vol. 6, pp. 85–92, Jun 2014.

[43] J. Wang, G. S. Pang, S. S. Chong, and C. G. Lee, “SNP web resources and their potential applications in personalized medicine,” Curr Drug Metab, vol. 13, pp. 978–90, Sep 1 2012.

[44] D. Gefel, I. Maslovsky, and J. Hillel, “[Application of single nucleotide polymorphisms (SNPs) for the detection of genes involved in the control of complex diseases],” Harefuah, vol. 147, pp. 449–54, 476, May 2008.

[45] D. A. Benson, M. Cavanaugh, K. Clark, I. Karsch-Mizrachi, D. J. Lipman, J. Ostell, et al., “GenBank,” Nucleic Acids Res, vol. 45, pp. D37–d42, Jan 4 2017.

[46] E. Capriotti and R. B. Altman, “Improving the prediction of disease-related variants using protein three-dimensional structure,” BMC Bioinformatics, vol. 12 Suppl 4, p. S3, 2011.

[47] Y. Choi, G. E. Sims, S. Murphy, J. R. Miller, and A. P. Chan, “Predicting the functional effect of amino acid substitutions and indels,” PLoS One, vol. 7, p. e46688, 2012.

[48] M. Hecht, Y. Bromberg, and B. Rost, “Better prediction of functional effects for sequence variants,” BMC Genomics, vol. 16 Suppl 8, p. S1, 2015.

[49] R. Calabrese, E. Capriotti, P. Fariselli, P. L. Martelli, and R. Casadio, “Functional annotations improve the predictive score of human disease-related mutations in proteins,” Hum Mutat, vol. 30, pp. 1237–44, Aug 2009.

[50] E. Capriotti, P. Fariselli, and R. Casadio, “I-Mutant2.0: predicting stability changes upon mutation from the protein sequence or structure,” Nucleic Acids Res, vol. 33, pp. W306–10, Jul 1 2005.

[51] J. Cheng, A. Randall, and P. Baldi, “Prediction of protein stability changes for single-site mutations using support vector machines,” Proteins, vol. 62, pp. 1125–32, Mar 1 2006.

[52] E. F. Pettersen, T. D. Goddard, C. C. Huang, G. S. Couch, D. M. Greenblatt, E. C. Meng, et al., “UCSF Chimera--a visualization system for exploratory research and analysis,” J Comput Chem, vol. 25, pp. 1605–12, Oct 2004.

[53] H. Ashkenazy, S. Abadi, E. Martz, O. Chay, I. Mayrose, T. Pupko, et al., “ConSurf 2016: an improved methodology to estimate and visualize evolutionary conservation in macromolecules,” Nucleic Acids Res, vol. 44, pp. W344–50, Jul 8 2016.

[54] B. Sun, Z. Huang, B. Wang, Y. Yu, S. Lin, L. Luo, et al., “Significance of Glypican-3 (GPC3) Expression in Hepatocellular Cancer Diagnosis,” Med Sci Monit, vol. 23, pp. 850–855, Feb 16 2017.

[55] C. Chen, X. Huang, Z. Ying, D. Wu, Y. Yu, X. Wang, et al., “Can glypican-3 be a disease-specific biomarker?,” Clin Transl Med, vol. 6, p. 18, Dec 2017.

[56] “UniProt: the universal protein knowledgebase,” Nucleic Acids Res, vol. 45, pp. D158–d169, Jan 4 2017.

[57] P. Mochalski, E. Diem, K. Unterkofler, A. Mundlein, H. Drexel, C. A. Mayhew, et al., “In vitro profiling of volatile organic compounds released by Simpson-Golabi-Behmel syndrome adipocytes,” J Chromatogr B Analyt Technol Biomed Life Sci, vol. 1104, pp. 256–261, Jan 1 2019.

[58] A. X. Zhu, P. J. Gold, A. B. El-Khoueiry, T. A. Abrams, H. Morikawa, N. Ohishi, et al., “First-in-man phase I study of GC33, a novel recombinant humanized antibody against glypican-3, in patients with advanced hepatocellular carcinoma,” Clin Cancer Res, vol. 19, pp. 920–8, Feb 15 2013.

[59] A. Das Bhowmik and A. Dalal, “Whole exome sequencing identifies a novel frameshift mutation in GPC3 gene in a patient with overgrowth syndrome,” Gene, vol. 572, pp. 303–6, Nov 10 2015.

[60] C. Kehrer, A. Hoischen, R. Menkhaus, E. Schwab, A. Muller, S. Kim, et al., “Whole exome sequencing and array-based molecular karyotyping as aids to prenatal diagnosis in fetuses with suspected Simpson-Golabi-Behmel syndrome,” Prenat Diagn, vol. 36, pp. 961–965, Oct 2016.

[61] Z. B. Alwi, “The Use of SNPs in Pharmacogenomics Studies,” Malays J Med Sci, vol. 12, pp. 4–12, Jul 2005.

